# Reference genomes of 545 silkworms enable high-throughput exploring genotype-phenotype relationships

**DOI:** 10.1101/2021.09.28.462073

**Authors:** Xiaoling Tong, Min-Jin Han, Kunpeng Lu, Shuaishuai Tai, Shubo Liang, Yucheng Liu, Hai Hu, Jianghong Shen, Anxing Long, Chengyu Zhan, Xin Ding, Qiang Gao, Bili Zhang, Duan Tan, Yajie Yuan, Nangkuo Guo, Yanhong Li, Zhangyan Wu, Lulu Liu, Chunlin Li, Yaru Lu, Tingting Gai, Yahui Zhang, Renkui Yang, Heying Qian, Yanqun Liu, Jiangwen Luo, Lu Zheng, Jinghou Lou, Yunwu Peng, Weidong Zuo, Jiangbo Song, Songzhen He, Songyuan Wu, Yunlong Zou, Lei Zhou, Linli Zhou, Lan Cheng, Yuxia Tang, Guotao Cheng, Lianwei Yuan, Weiming He, Jiabao Xu, Tao Fu, Yang Xiao, Ting Lei, Anying Xu, Ye Yin, Jian Wang, Antónia Monteiro, Eric Westhof, Cheng Lu, Zhixi Tian, Wen Wang, Zhonghuai Xiang, Fangyin Dai

**Affiliations:** State Key Laboratory of Silkworm Genome Biology, Institute of Sericulture and Systems Biology, Southwest University, Chongqing 400715, China; Key Laboratory of Sericultural Biology and Genetic Breeding, Ministry of Agriculture and Rural Affairs, College of Sericulture, Textile and Biomass Sciences, Southwest University, Chongqing 400715, China; BGI Genomics, BGI-Shenzhen, Shenzhen 518083, China; State Key Laboratory of Plant Cell and Chromosome Engineering, Institute of Genetics and Developmental Biology, Innovation Academy for Seed Design, Chinese Academy of Sciences, Beijing 100101, China; Chongqing Sericulture Science and Technology Research Institute, Chongqing 400715, China; Jiangsu Key Laboratory of Sericulture Biology and Biotechnology, School of Biotechnology, Jiangsu University of Science and Technology, Zhenjiang, Jiangsu 212018, China; College of Bioscience and Biotechnology, Shenyang Agricultural University, Shenyang, Liaoning 111000, China; Shaanxi Key Laboratory of Sericulture, Ankang University, Ankang, Shaanxi 710072, China; Institute of Sericulture and Agricultural Products Processing, Guangdong Academy of Agricultural Sciences, Guangzhou, Guangdong 510000, China; BGI-Shenzhen, Shenzhen 518083, China; James D. Watson Institute of Genome Sciences, Hangzhou 310058, China; Biological Sciences, National University of Singapore, 14 Science Drive 4, Singapore 117543, Singapore; Science Division, Yale-NUS College, Singapore 138614, Singapore; Architecture et Réactivité de l’ARN, Institut de Biologie Moléculaire et Cellulaire, UPR9002 CNRS, Université de Strasbourg, Strasbourg 67084, France; University of Chinese Academy of Sciences, Beijing 100049, China; School of Ecology and Environment, Northwestern Polytechnical University, Xi’an, Shaanxi 710072, China; Kunming Institute of Zoology, Chinese Academy of Sciences, Kunming, Yunnan 650204, China

## Abstract

The silkworm Bombyx *mori* is a domestic insect for silk production and a lepidopteran model. The currently available genomes limit a full understanding of its genetic and phenotypic diversity. Here we assembled long-read genomes of 545 domestic and wild silkworms and constructed a high-resolution pan-genome dataset. We found that the silkworm population harbors extremely variable genomes containing 7,308 new gene families, 4,260 (22%) core gene families, and 3,432,266 non-redundant SVs. We deciphered a series of causal genes and variants associated with domestication, breeding, and ecological adaptation traits, and experimentally validated two of those genes using CRISPR-Cas9 or RNA interference. This unprecedented large-scale genomic resource allows for high-throughput screening of interesting traits for functional genomic research and breeding improvement of silkworms and may serve as a guideline for traits decoding in other species.

## Introduction

Long-read genomes at population scale in species facilitate insights into the intra-species genomic contents and the contributions of structure variations (SVs) in traits determination^1-3^. Recently, increasing pan-genomes generated from tens of samples in a species have revealed that SVs have extensive distribution in the genome, high diversity among individuals, and significant contribution to phenotype determination^1, 2, 4-13^. Pan-SVs present an “open” growth curve without a plateau^5, 14^. Thus, a very large number of representative high-quality genomes would be needed to comprehensively understand the entire genetic variation of a species.

Silkworm, *Bombyx mori*, is an important economic insect producing silk known as the “queen of fabrics”. It is raised in 58 countries^15^ and generates a global trade volume above US$50 billion annually. Silkworm is also an ideal model insect for scientific research, including genetics, evolution, physiology, genomics, and genetic engineering^16-20^. Owing to long-term artificial and natural selection, numerous silkworm resources exhibiting extensive phenotypic variations were preserved worldwide (Extended Data Fig. 1), including local strains, improved varieties, genetic stocks (spontaneous and induced mutants), and wild silkworms. However, the currently available genomes^21, 22^ limited mining and utilization of these precious resources. To address this issue, we generated a large-scale of long-read assemblies from 545 silkworms for underpinning the basic and applicational research.

## Results

### Deep re-sequencing of 1,078 silkworms

To explore the full genomic diversity within silkworm species, we collected silkworms (1,078 samples) as comprehensively as possible, containing 205 local strains, 194 improved varieties, 632 genetic stocks, and 47 wild silkworms (Fig. 1a, Supplementary Table 1). A total of 31.52 Tb next-generation sequencing (NGS) reads with an average coverage depth of ∼65× for all the samples were obtained (Supplementary Table 1). Using NGS data of our 1,078 silkworms and four previously released wild silkworm genomes, 43,012,261 single nucleotide polymorphisms (SNP) and 9,344,375 small insertions or deletions (Indel, < 50 bp) were identified. The SNP and Indel density were one SNP per 11 bp and one Indel per 49 bp, indicating silkworm has a high degree of genomic diversity.

**Fig. 1.**
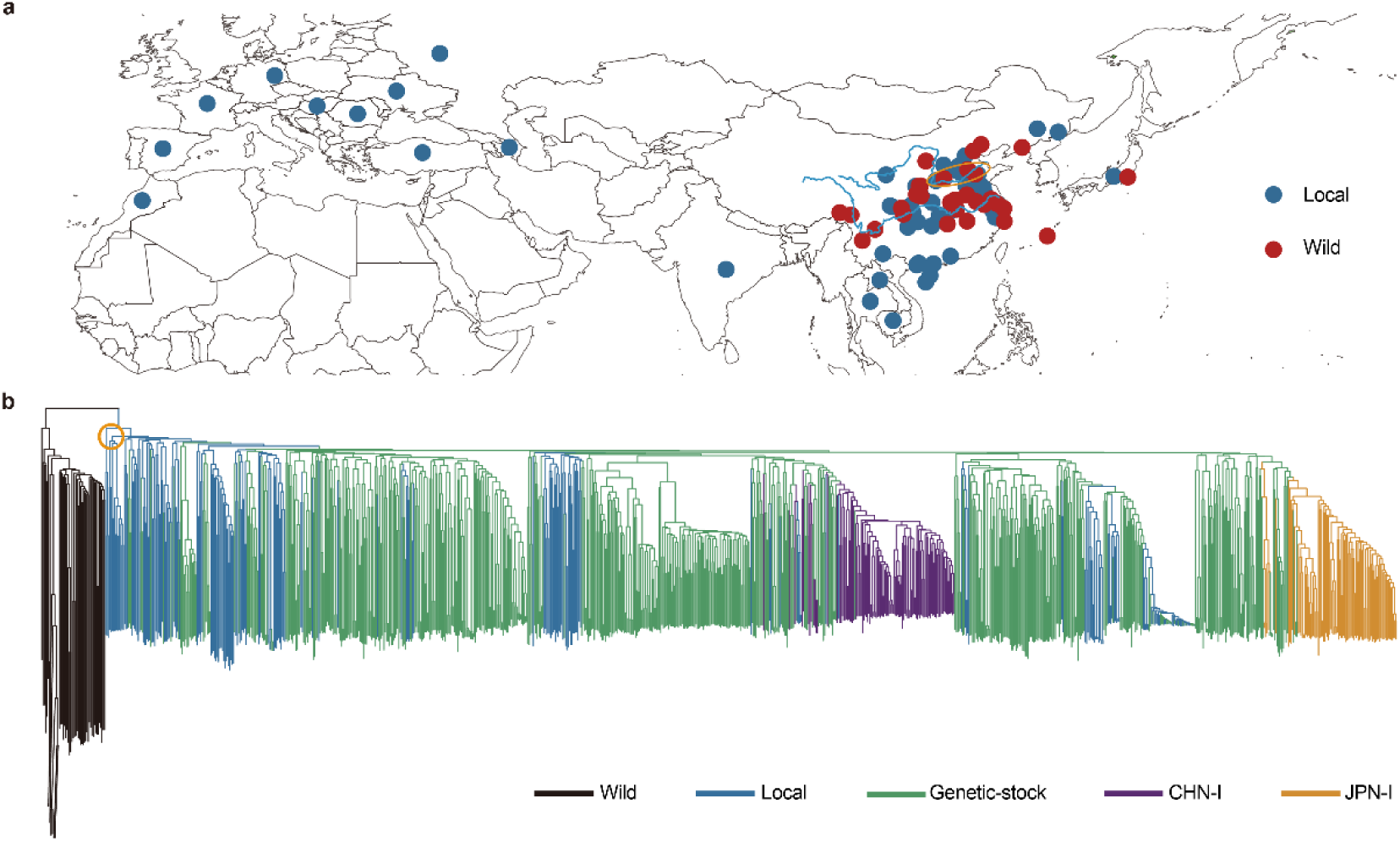
Geographic distribution and phylogenetic tree of silkworm. **a**, Geographic distribution of 1,082 silkworms. The blue and red dots represent the local and wild silkworms. The orange ellipse marks the middle and lower reaches of the Yellow River area. **b**, Phylogenetic tree based on genome-wide SNPs of 1,082 samples. The clades in the orange circle represent the local strains from the middle and lower reaches of the Yellow River area.

Two interesting findings were revealed from the phylogenetic analysis of these 1,082 genomes. First, our result resolved a long-term controversy about the origins of domestic silkworms, either from single or multiple geographical sites^23-25^. The local strains from the middle and lower reaches of the Yellow River are placed at the basal position of the domestic silkworm clade (Fig. 1b), suggesting that silkworms were domesticated from a single geographical site in China. Second, the highly concentrated distribution of improved strains in two subgroups reflects an extremely narrow genetic base of the commercial silkworms (Supplementary Table 2). They were probably descended from a limited ancestral genetic resource in China (CHN-I) and Japan (JPN-I) independently. It is essential to exploit and utilize the abundant genetic resource for future silkworm breeding.

### Long-read genomes of 545 silkworms

To reach an overview of genomic content in silkworms, 545 representatives from each group evenly covering the phylogenetic tree were selected to perform long-read sequencing and genome assembly (Fig. 2a, Extended Data Fig. 2a). A total of 24.06 Tb third-generation sequencing (TGS) read data with an average sequencing depth of 98× were obtained (Fig. 2b) and an average read length of 23.5 kb (Fig. 2c). *De novo* assemblies of these 545 genomes showed that the average genome size is 457.9 Mb with an average contig N50 size of 7.6 Mb (Fig. 2d) and an average repeat sequence proportion of 51% (Fig. 2e). The BUSCO evaluation value and mapping ratio of NGS reads to the assembled genomes is 98% and 99% on average (Fig. 2f, Supplementary Table 3), indicating that the assembled genomes have high completeness.

**Fig. 2.**
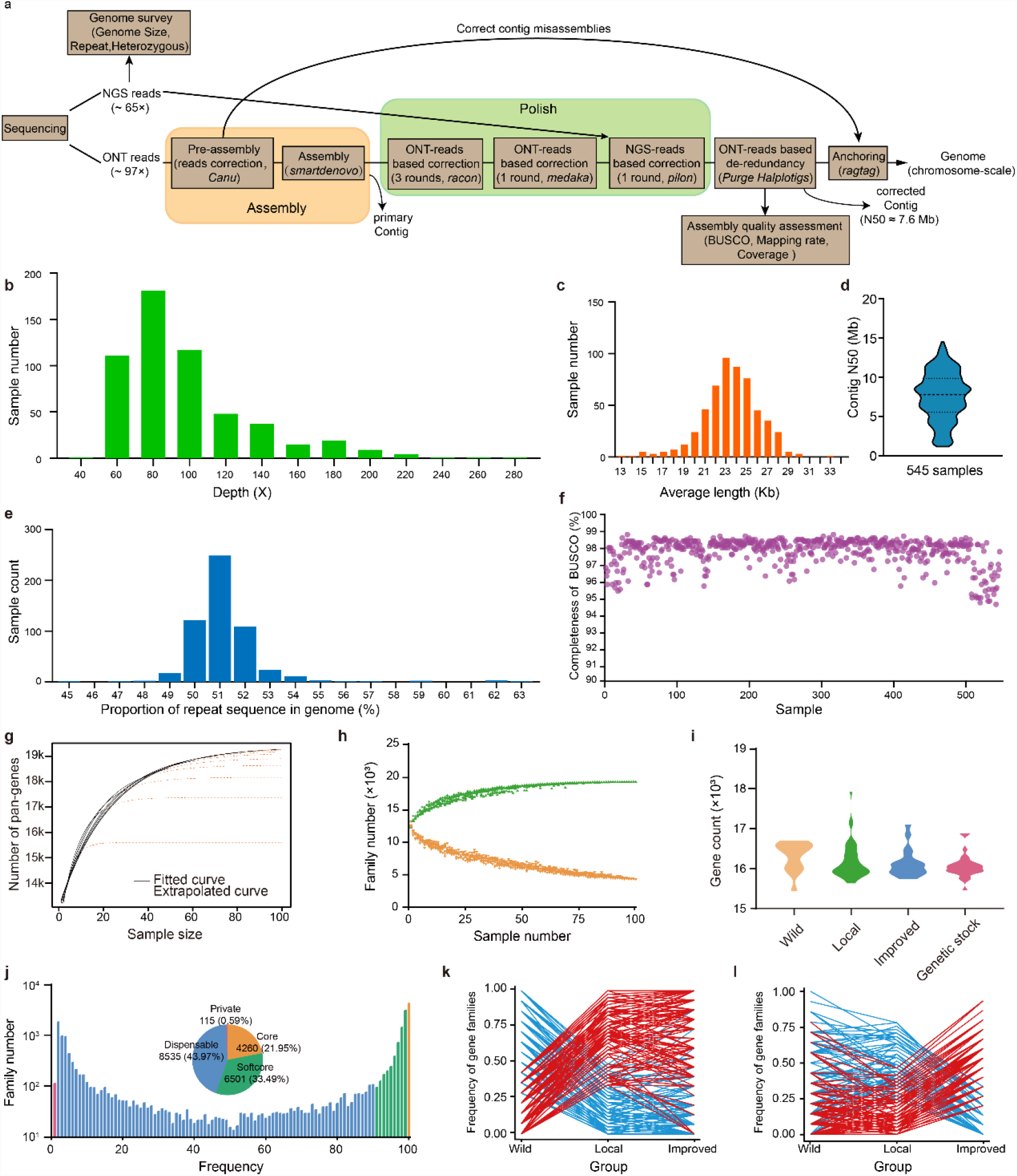
Sequencing, assembly, and pan-gene analysis of silkworm genomes. **a**, The strategy of genome sequencing and assembling. **b**, The average read coverage distribution of 545 strains. **c**, The average read length distribution of 545 strains. **d**, The distribution of contig N50 length of the 545 genomes. **e**, The distribution of repeat sequences proportion in the 545 genomes. **f**, The BUSCO evaluation values of the 545 genomes. **g**, Evaluation of pan-gene plateau. The black curves are fitted with actual data, and the yellow dotted curves are extrapolated by the model of *y* = *A* + *Be*^*Cx*^. The pan-genes obtained from 80, 90, and 100 genomes are similar. **h**, Counts of pan-gene (green) and core-gene (orange) families with increased samples. **i**, There is no significant difference in gene counts among wild, local, improved, and genetic stock groups. **j**, The histogram shows gene family number with different frequencies. The pie chart shows the proportion of core, softcore, dispensable, and private genes in those genomes. **k** and **l**, gene families with significantly frequency differentiation in wild-local comparison (k) and local-improved comparison (l). In the two comparisons, gene families with significantly increased (red) or reduced (blue) frequencies in domestication (k) and improvement (l) were showed with different colour.

To showcase full landscape of silkworm genes, we first estimated how many genomes are enough to capture the full set of genes of silkworms using a gradually superimposed approach, and finally annotated 100 genomes for which pan-genes reached a plateau (Fig. 2g, h, Extended Data Fig. 2b). These genomes contain an average of 16,234 genes (Fig. 2i, Supplementary Table 4). A total of 19,411 gene families were identified in the 100 genomes, containing 4,260 (22%) core (shared by all 100 samples), 6,501 (34%) softcore (shared by > 90% samples but not all), 8,535 (44%) dispensable (shared by more than one but ≤ 90% samples), and 115 (1%) private (present in only one sample), respectively (Fig. 2j, Extended Data Fig. 2c). Among the four categories of genes, core genes are supposed to be more conserved in functions and may play important roles in gene regulation based on the results of functional annotation, expression, and *d*_*N*_/*d*_*S*_ analyses (Extended Data Fig. 2d-j).

In these 19,411 gene families, 7,308 (38%) families are new genes absent in the prior reference genome. Around 83% (5,807) of the novel gene families have GO items, or transcriptional evidence (Supplementary Table 4), and ∼99% of the new families present in more than two genomes (Extended Data Fig. 2i), suggesting that they are truly present and informative for further functional genomics studies in silkworm. Distribution analysis showed that 251 and 241 gene families had a significant frequency change (FDR < 0.0001 and fold change > 2) in wild-local and local-improved population comparisons, respectively (Fig. 2k, l). Among which, 72% and 82% were novel families, indicating crucial roles of new identified genes for silkworm domestication and breeding, further, for differentiation of subpopulation.

### SV characters and graph-based pan-genome

To construct a high-resolution silkworm pan-genome, we mapped the long reads of each of the 545 genomes to the reference genome^22^ and obtained an average of 120,216 SVs per genome (Extended Data Fig. 3a). The average SV count in wild genomes is significantly higher (*p* < 0.0001, *t*-test) than in domestic silkworms (Fig. 3a). Insertions (INS) and deletions (DEL) (referred as PAVs, presence/absence variations) constitute most of the SVs (Fig. 3b, Supplementary Table 3). Further, all SVs were merged to generate 3,432,266 non-redundant SVs (nrSVs) with a majority (96%) shorter than 15 kb length and a large proportion (81%) of rare alleles (allele frequencies less than 0.05) (Fig. 3c, d), presenting an open silkworm pan-SVs (Fig. 3e, Extended Data Fig. 3b) and a high SV density (one SV per 134 bp) in the silkworm population (Fig. 4a).

**Fig. 3.**
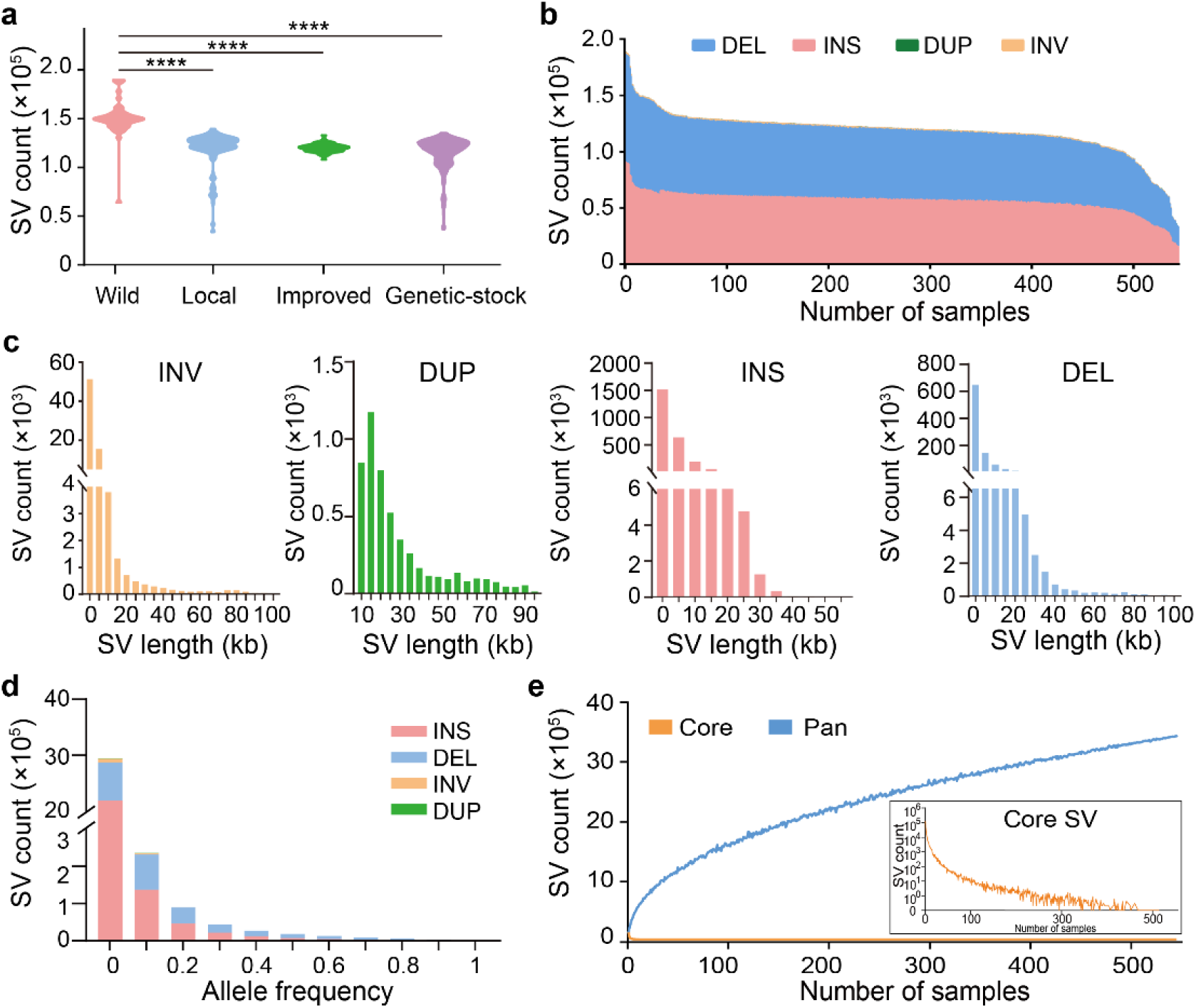
Characterization of pan-SVs in 545 silkworm genomes. **a**, Violin plots of SV counts in the four genetic resource groups. **b**, SV count of insertions (INS), deletions (DEL), duplications (DUP), and inversions (INV) in each of the 545 genomes. The proportions of DUP and INV are too low to be observed in the graph. **c**, The distribution of insertion (INS), deletion (DEL), inversion (INV), and duplication (DUP) length. **d**, Allele frequency of nrSVs from 545 samples. **e**, Pan-SV and core-SV counts with additional genomes.

**Fig. 4.**
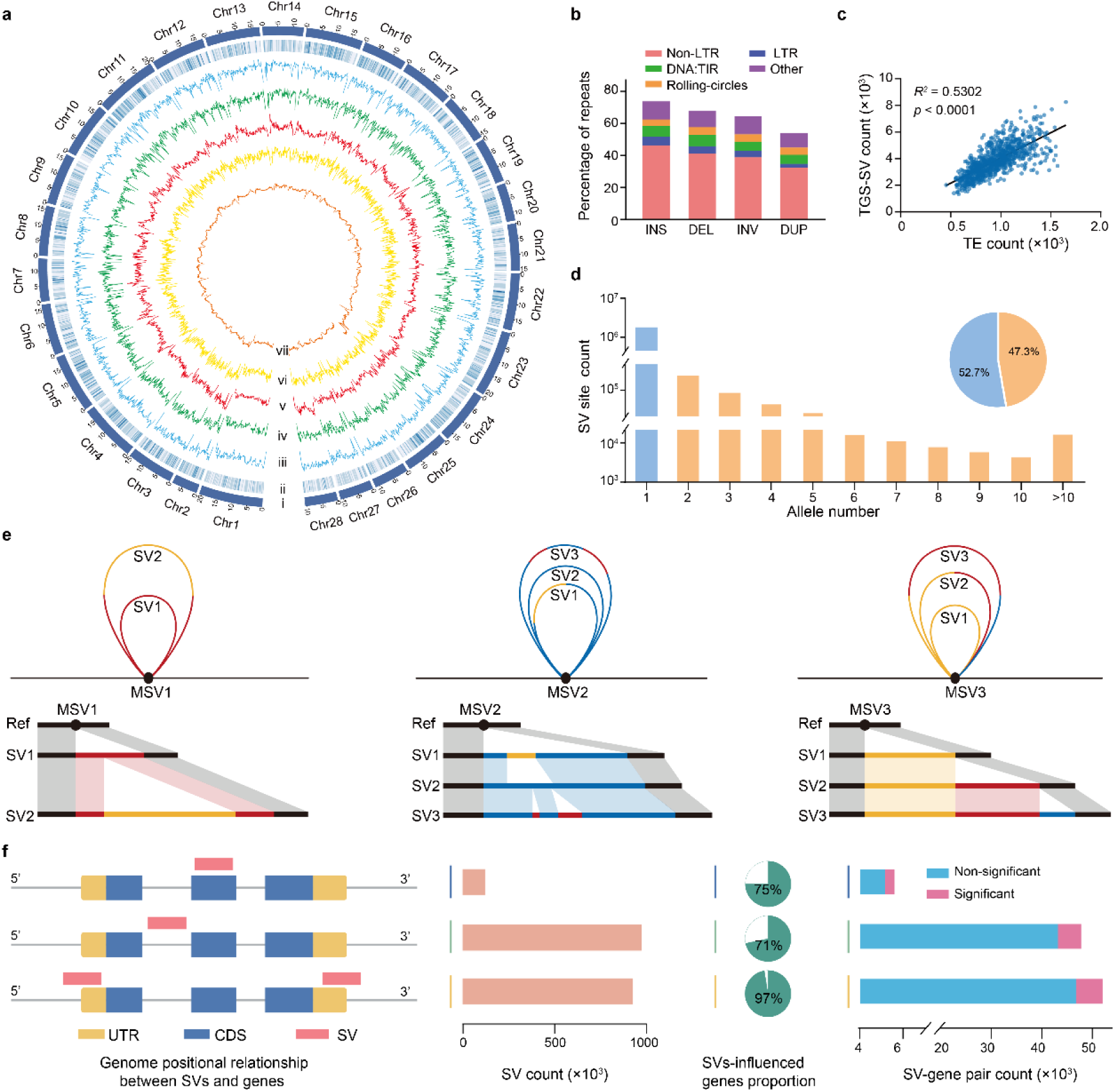
Characterization of non-redundant SVs (nrSVs). **a**, Distribution map of genetic variations in 1,082 genomes. (i) Chromosomes. (ii) Gene density. (iii) SNP and (iv) Indel densities across 1,082 genomes. (v) Non-redundant SVs density of 545 TGS genomes. (vi) Density of non-redundant SVs of 537 NGS-only genomes. (vii) TE density. **b**, Components of transposable elements (TEs) in sequences of insertion (INS), deletion (DEL), inversion (INV), and duplication (DUP). **c**, Correlations of TGS-SV and TE counts. TGS-SV and TE numbers were counted in uninterrupted 500 kb windows. There is a significant linear relationship between SVs and TEs distribution on chromosomes (*R*^*2*^ = 0.53, p < 0.0001). **d**, A single or multiple SV (MSV, ranging from 2 to 135) alleles were found in a certain genomic site. Histogram shows SV site counts with different allele numbers. Pie shows proportions of SV site with single (blue) and multiple (yellow) alleles. **e**, Schematic diagrams of genomic sites with MSV alleles and their collinearity. **f**, SVs impact on genes. SV count (middle) in the corresponding region (left) and their impact on gene expression (right). The pie charts show the proportion of genes influenced by SVs.

The average SV count per genome in silkworms, to our knowledge, is 4-5 times that of previously examined eukaryotes^5, 8, 12, 13^. A primary explanation could be that silkworm genome harbors a higher density (2,075 copies per Mb per genome) of transposable elements (TEs) compared with these species (229-1,222 copies per Mb per genome), since TE is regarded as the major contributor to genome SVs. Indeed, we found that TEs constitute the largest component (67%) of SVs sequences in silkworm genomes (Fig. 4b). Further, the distribution of SVs on chromosomes is significantly related to that of TEs (*R*^2^ = 0.53, *p* < 0.0001) (Fig. 4c). Another possible explanation could be that SVs events occurred with a high mutation rate, reflected by the appearance of a high proportion (47%) of multiple SVs (MSVs, ranging from 2 to 135 nrSVs in a single site) alleles in all the silkworms (Fig. 4d, e) and a large proportion (71%) of SVs in domestic silkworms occurred after domestication. Moreover, domestic silkworms are more tolerant to deleterious or slightly deleterious mutations (rare alleles) because they are entirely dependent on humans and under weak natural selection pressure. This speculation is supported by our finding that the proportion of rare allele SVs in domestic silkworm (70%) is higher than in wild silkworm (56%).

All PAVs were integrated into the linear reference genome^22^ to construct a graph-based pan-genome. Mapping the short reads of the remaining 537 NGS sequenced genomes to the pan-genome led to the identification of 59,037 novel SVs for a total of 454,671 SVs (Fig. 4a). The distribution of SVs on chromosomes called by NGS data is like that of SVs identified by TGS data (Extended Data Fig. 3c, d). It suggests that the pan-genome could be used as a comprehensive reference to analyze genomic variations in the short-read sequenced genomes.

### SVs impact on genes

To investigate the effects of SVs on genes, we analyzed the relative genomic positions of SVs. We found that 98% (16,460) of reference genes harbor SVs (55% of total SVs) in their potential expression regulatory regions (PERRs, including the introns and ±5 kb flanking regions of a gene, in this study) or coding sequence (CDS), indicating a large impact of SVs on gene expression and structure. Among these SVs, ∼93% (1,762,169) are detected in PERRs of 16,456 genes (97% of reference gene) (Fig. 4f). In the seven strains on which RNA-seq was performed, 78,797 SVs were found in PERRs of 14,796 genes and 99,875 SV-gene pairs were constructed. Finally, 4,402 genes in 9,417 SV-gene pairs were found to be differentially expressed (FDR < 0.001) in at least one tissue between strains with and without corresponding SV (Fig. 4f). The other 7% (130,669) SVs influence the CDS of 12,661 genes (75% of reference gene). In contrast, 5,637 and 9,527 genes are influenced respectively by SNPs and Indels (< 50 bp) at the sites of gene structure determination (e.g. introducing a new stop codon, or leading to a frameshift) (Supplementary Table 5), which implies a more extensive impact of SVs than Indels and SNPs on protein coding genes.

### SVs contribute largely to silkworm domestication

In order to identify genomic loci related to silkworm domestication, we examined two signatures of strong selection in genomes: changes in allele frequencies (AFs) as well as areas of depleted genetic variation. We identified 5,354 dSVs (domestication-associated SVs) which potentially played roles in domestication of silkworm because their AFs show differences (FDR < 0.0001, fold change > 2) in wild and local silkworms (Fig. 5a). Moreover, we identified 395 (2.3% of the whole-genome genes) domestication-associated genes (DAGs) based on the selective sweep analysis using SNPs. A total of 690 dSVs were identified in or close to 57% (225) of the DAGs (Supplementary Table 6), suggesting a broad impact of SVs on silkworm domestication.

**Fig. 5.**
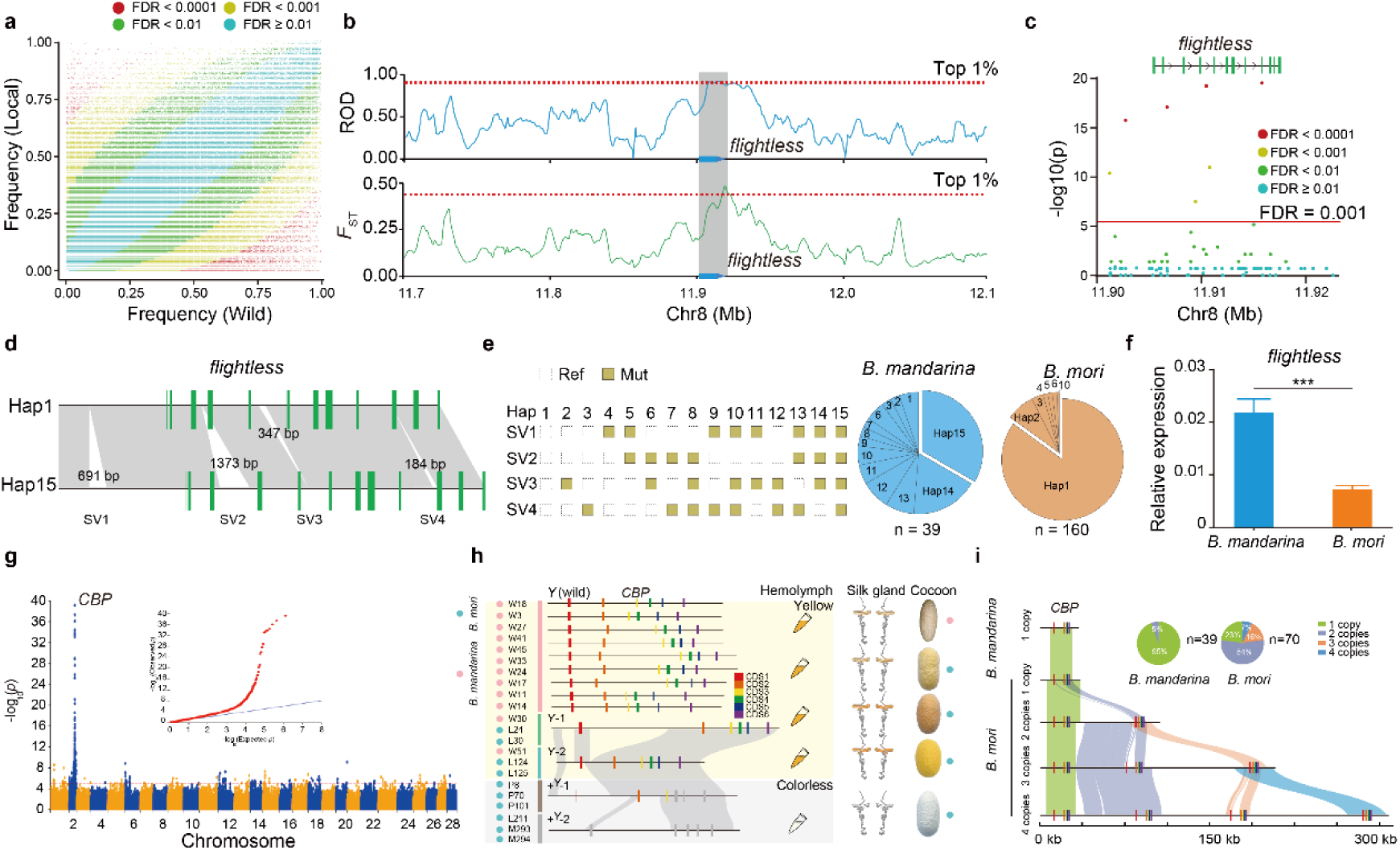
SVs associated with domestication. **a**, Frequency distribution of SVs in wild and local groups, dots represent SVs. **b**, Selective sweep signatures of *flightless. F*_ST_ and ROD (reduction of diversity) between wild and white domestic silkworms are shown. Red dashed lines represent the threshold value of top 1% selection signatures. **c**, SV frequency divergence between wild and domestic silkworms on *flightless*, and their flanking ±5 kb. **d**, The frequency divergence of SVs in *flightless* and its flanking ±5 kb between wild and local silkworms. **e**, Haplotypes of SVs within *flightless*. Pie charts show haplotype proportions of *flightless* in *B. mandarina* and *B. mori*. **f**, Differential expression of *flightless* in the flight muscle of *B. mandarina* and *B. mori*. ***, *p* < 0.001, *t*-test. **g**, SV-based GWAS of hemolymph color (yellow and colorless) in domestic silkworms. **h**, The schematic diagram shows the gene structures and collinearity relationships of different *CBP* types. The gray rectangles represent lost CDS in the *+*^*Y*^-1 and *+*^*Y*^-2. **i**, Collinearity and proportion of *CBP* gene with different copy numbers in domestic and wild silkworms.

The degeneration of flight capacity in *B. mori* is regarded as a typical consequence of the “domestication syndrome” (DS) but its genetic basis is unknown. A selective sweep analysis showed that a signature of positive selection can be detected in the gene body and regulatory regions of *flightless* (*flil*, a flight muscle structural gene) (Fig. 5b). Four SVs with significant frequency divergence between wild and domestic silkworms were found in the regulatory regions of *flil* and formed 15 haplotypes (Fig. 5c-e). Hap1 lacks these 4 SVs and is present in 85% of *B. mori* samples (Fig. 5e). Further, the expression of *flil* is significantly lower in *B. mori* than in *B. mandarina* (Fig. 5f), which might be influenced by the SVs. In our prior study, degeneration of flight muscle was found in *B. mori*^26^. Correspondingly, in *Drosophila*, loss of function of *flil* makes adults unable to fly^27^. We suggest, therefore, that the SVs influence the expression of *flil* and contributed to flight degeneration in silkworm.

Natural variation in the color of cocoons has been exploited in sericulture but the genetic basis of the variation has been poorly explored. Generally, *B. mandarina* has yellow-colored hemolymph (*Y*) and yellowish cocoons due to carotenoids accumulation. Humans selected white cocoons for better dyeing qualities while brilliant yellow-red cocoons for their eco-friendly character and natural functional pigments. The CBP, a transporter of carotenoids crosses the midgut and the silk gland^28^, is strongly associated with the artificial selection of hemolymph color, influencing subsequent cocoon color. An SV-based genome-wide association study (GWAS) on hemolymph color identified a region with a significant signal that contains *CBP* (Fig. 5g). In *B. mandarina*, 95% of samples have only one copy of functional *CBP* (*Y*) with varied introns (Fig. 5h). In *B. mori*, two *CBP* types (*Y*-1 and *Y*-2) derived from wild silkworm were observed in the population producing yellow hemolymph. While two null *CBP* mutations (*+*^*Y*^-1 and *+*^*Y*^-2) originated from *Y*-2 were identified in the colorless hemolymph strains (Fig. 5h, Extended Data Fig. 4). Significantly, in domestic silkworms, a majority (77%) of yellow blood strains contain 2-4 *CBP* copies (Fig. 5i). In contrast, colorless blood strains harbor only one mutated *CBP* copy. The results reveal the complex evolutionary paths of *CBP*, which has undergone gene duplication or gene loss for differential selection aims.

### Genetic basis of breeding

To identify genes and SVs associated with silkworm breeding, we performed comparative analyses among improved and local strains. CHN-I and JPN-I are the improved groups currently used for generating heterosis (or hybrid vigor). We identified 246 (CHN-I) and 228 (JPN-I) improvement associated regions (IARs) containing 281 improvement associated genes (IAGs) in each group (Fig. 6a). Interestingly, the two improved groups only shared less than 10% of the IARs (containing 51 common genes) with occasional intergroup introgression (Extended Data Fig. 5a, b), suggesting that breeding proceeded largely independently in each group, and revealing the genetic bases of silkworm heterosis. We found 261 iSVs (improvement-associated SVs, showing significant divergent frequencies between improved and local groups) in the gene and flanking region (± 5 kb) of 111 IAGs (Extended Data Fig. 5c, d, Supplementary Table 6). More than 98% of these iSVs are within the PERRs of these genes, suggesting that selection in PERR plays a major role in silkworm breeding.

**Fig. 6.**
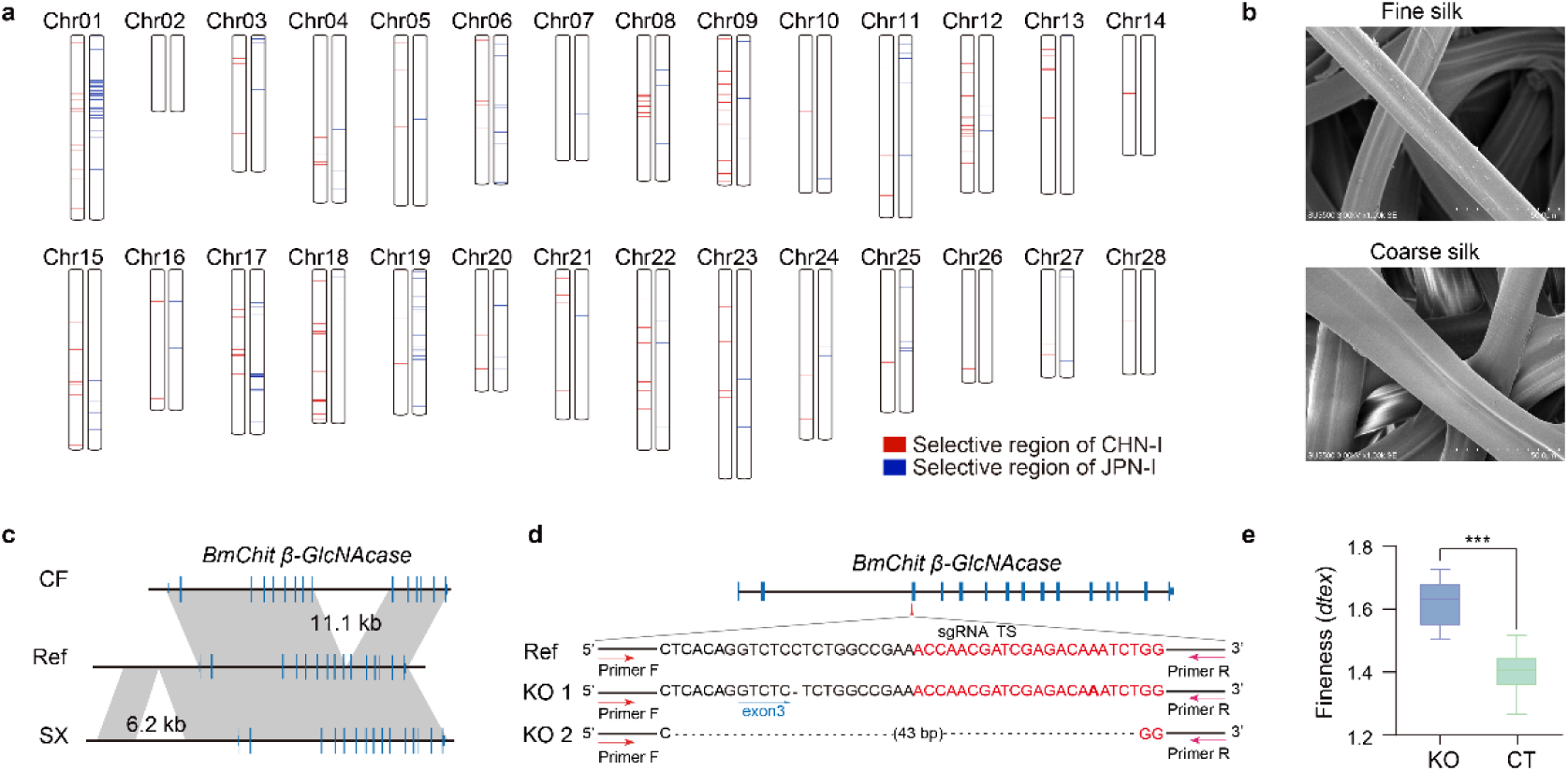
Genetic basis of heterosis and fiber fineness. **a**, Selective regions of CHN-I (red, Chinese improved strains) and JPN-I (blue, Japanese improved strains) in process of breeding. **b**, Silk with fine and coarse silk under scanning electron microscope. bar = 50 μm. **c**, An 11.1 kb insertion in the intron of *BmChit β-GlcNAcase* in CF strain and a 6.2 kb insertion downstream from *BmChit β-GlcNAcase* in SX strain. **d** and **e**, CRISPR-Cas9 mediated *BmChit β-GlcNAcase* knockout (KO) generated two types of mutations (d) and coarse silk fineness compare to control (CT) strain (e). sgRNA target site (TS) marked in red. ***, *p* < 0.001, Student’s *t*-test.

Fine silk has higher economic value in sericulture but the genetic basis of fiber fineness is unknown. We previously found that a part of the spinneret, the silk press, is related to fineness through comparisons between fine silk (SX, CF) and coarse silk (XF, QB) strains^29^ (Fig. 6b, Extended Data Fig. 5e). Here we performed RNA-seq of the silk press in these four strains and identified 40 differential expressed genes (DEGs) (Extended Data Fig. 5f). By scanning for variations in the genomic regions of these DEGs, we identified an 11.1 kb insertion in intron and a 6.2 kb downstream insertion of *chitooligosaccharidolytic beta-N-acetylglucosaminidase* (*BmChit β-GlcNAcase*) gene in CF and SX strains, respectively (Fig. 6c). *BmChit β-GlcNAcase* was expressed significantly more in fine silk strains (SX, CF) and had an expression peak in silk press at the wandering stage (a stage at the start of spinning) (Extended Data Fig. 5g, h). CRISPR-cas9 mediated knockout of *BmChit β-GlcNAcase* produced increased fineness (Fig. 6d, e). All these results suggest a key role of *BmChit β-GlcNAcase* in silk fineness determination.

### Decoding insect adaptation using the pan-genome

The silkworm moltinism (the number of larval molts) exhibit obvious phenotypic variability along environmental gradients. For instance, we found that most tetramoulting (*M*^4^) (∼94%) silkworms are from tropical regions. In contrast, most trimoulting (*M*^3^) (∼70%) silkworms come from northern China (Fig. 7a). To identify moltinism-associated genes, we first compared *M*^3^ with *M*^4^ genomes and searched for divergent regions based on SNPs. The result revealed that the top divergent signatures are located in the region of 5.36-5.55 Mb of chromosome 6, with only a *BmScr* gene present (Fig. 7b). Previous studies suggested that *BmScr*, a homeodomain-containing transcription factor, could affect moltinism through regulating juvenile hormone (JH) biosynthesis^30^. Furthermore, the result of SVs-based GWAS of *M*^3^ and *M*^4^ lines revealed a significant signal on a 150 bp PAV in the first intron of *BmScr* (Fig. 7c, d), indicating this PAV could be one major causal mutation for moltinism variation in silkworms. The present study, at the population level, reveals for the first time that *BmScr* is related to silkworm environmental adaptation (moltinism).

**Fig. 7.**
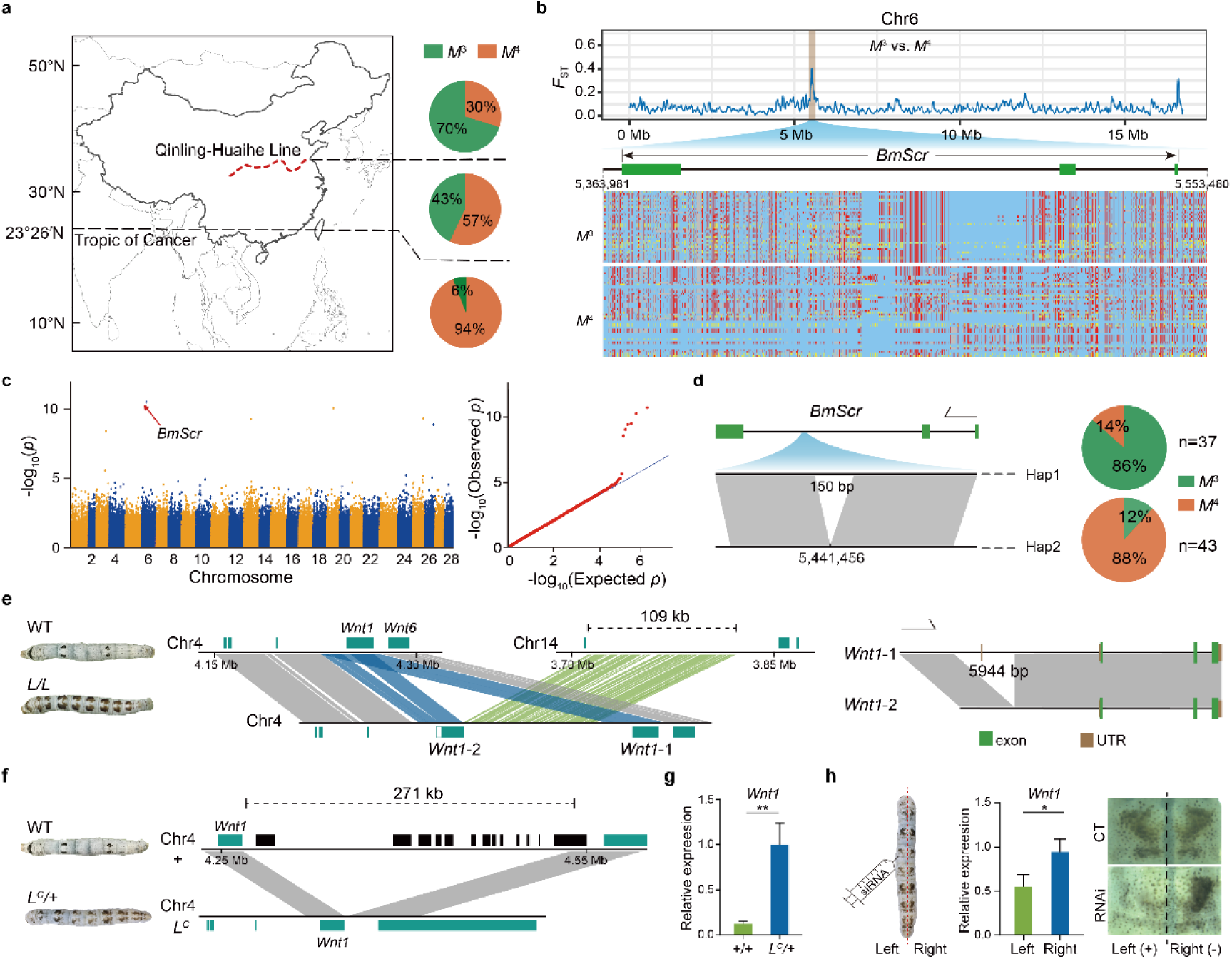
Silkworm moltinism and aposematic coloration pattern. **a**, Pie charts show the distributions of local silkworms with different type of moltinism (*M*^3^ and *M*^4^) in three geographic regions divided by Qinling Mountains-Huaihe River Line (Qinling-Huaihe Line) and Tropic of Cancer. **b**, Line charts show population divergence (*F*_ST_) between *M*^3^ and *M*^4^ on Chromosomes 6. Haplotype view of SNPs in *M*^3^ and *M*^4^ in regions with higher *F*_ST_ value on chromosomes 6. **c**, SV-based GWAS of moltinism (*M*^3^ and *M*^4^). The red arrow represents a significant signature (a 150 bp SV) in *BmScr*. **d**, Two haplotypes of *BmScr* gene defined by the presence or absence of the 150 bp in the second intron. Pie charts show the proportion of *M*^3^ and *M*^4^ strains within each haplotype. **e** and **f**, Compared to wild type (WT), large insertion (green, 109 kb fragment derived from chromosome 14) and duplication (blue, duplication of *Wnt1* and its flanking sequences) were identified in *L* strains (e), and a 271 kb deletion was identified in *L*^*C*^ (f). **g**, The expression of *Wnt1* is significantly increased in heterozygous *L*^*C*^. **h**, The caltrop-type marking was lost and the *Wnt1* expression of the left side (*Wnt1* RNAi) was significantly decreased compared to right side (control) after *Wnt1* knockdown. **, *p* < 0.01, *, *p* < 0.05, Student’s *t*-test.

Aposematic coloration, in the form of conspicuous body markings, is another important adaptive trait. The silkworm alleles *Multilunar* (*L*) and *Caltrop-type multilunar* (*L*^*C*^) lead to similar twin-spot markings on the dorsal side of the larval body (Fig. 7e, f), which are commonly used as aposematic markings to avoid predators in insects^31, 32^. A prior study suggested that the *L* phenotype is caused by short sequence changes without large variations in the 5’-flanking 19-kb region of *Wnt1* on chromosome 4^33^. Here, we identified two *L* specific large SVs, a duplication (34 kb) containing an additional *Wnt1* copy (named *Wnt1-2*) with a 5,944 bp deletion in its cis-regulatory region and an insertion (109 kb) derived from chromosome 14 in the 3’ terminal 500 bp site of *Wnt1-2* (Fig. 7e). We thus speculated that the *L* phenotype was caused by those large SVs rather than by short sequence changes. In addition, we found an *L*^*C*^ specific large deletion (271 kb) in the 3’ flanking region of *Wnt1* (Fig. 7f). The *Wnt1* expression in the epidermis of heterozygous *L*^*C*^ (*L*^*C*^/+) is significantly higher than that in normal strains (+/+) (Fig. 7g). *Wnt1* RNAi in the left side of the epidermis of *L*^*C*^ (*L*^*C*^/+) larvae blocked spot marking formation (Fig. 7h). These results reveal that large and complex SVs in *L* alleles, which cannot be obtained by mapping cloning, affect the expression pattern of *Wnt1* and result in twin-spot markings variation.

## Discussion

To our knowledge, this project has sequenced the largest number of long-read genomes in a eukaryotic species except human. These high-quality genomes have provided unprecedented resources for comprehensively understanding of silkworm genomic variations and phenotypic diversity. In the present study, we decoded the causal genes and variants of several important traits related to the “domestication syndrome” (flightlessness), breeding (silk fineness and cocoon color), and ecological adaptation (moltinism, and aposematic coloration). Furthermore, 632 well-described genetic stocks were used in our study. They are valuable genetic resources exhibiting various morphological, biochemical, and behavioral traits affecting different metamorphic stages. Accessing to the full genomes of these genetic resources will benefit insect science. Our findings have important significance for silkworm breeding improvement and general science. Importantly, these results show that the large-scale high-quality genomes of a certain species are powerful in traits decoding, which has general implications for understanding of phenotypic diversity of other species.

In summary, our results show that the pan-genome based on hundreds of long-read sequenced genomes provided unprecedented resources for high-throughput and accurate assessment of genotype-phenotype associations. This ushers a new era for silkworm basic research, a new generation of molecular breeding, and provides new guidelines for insect science, pest control, and traits decoding in other species.

## Methods

### Silkworm collection

We collected 1078 silkworms, containing 205 local strains from diverse geographic regions of the traditional silk producing countries and adapted to different environmental conditions (e.g., China, Japan, Korea, India, Thailand, Laos, Vietnam, Russia, France, Italy, Germany, Hungary, Spain, Turkey, Romania, Morocco, Cambodia, Azerbaijan, Ukraine, and Bulgaria), 194 improved varieties showing desirable properties for commercial breeding (e.g., high yield and quality silk, natural color cocoon, greater robustness, uniform development, and high hatchability), 632 genetic stocks (natural or artificial mutants), and 47 wild silkworms from a full range of their geographic distributions in China. It is worth mentioning that the silkworm genetic stocks collected from all over the world and maintained since the 1900s constitute a unique resource for insect biology and specifically for Lepidoptera. They harbor diverse characteristics such as embryonic lethality, colors and shapes of the eggs, colors and patterns of the larval cuticle and the adult wings, body shapes, colors and shapes of the cocoon, abnormal synthesis of certain protein (such as sericin and fibroin), and physiological defects in reproductive system. More than 400 mutants have been mapped to all of the 28 linkage groups of silkworms using classical linkage analysis^34^.

Most collections (∼90%) are from the silkworm genetic resource banks in Southwest University (Chongqing, China). A minority of germplasms was collected from other universities and sericulture research institutes.

### Short-read sequencing

For next-generation sequencing (NGS), a paired-end sequencing library of each sample of 1,078 silkworms was constructed with insertion sizes ranged from 300 to 400 bp and sequenced through DNBSEQ platform of BGI-Shenzhen. *SOAPnuke* v1.5.6^35^ was used to filter out the low-quality reads (a read containing over 40% low-quality bases, base quality value less than 20 was considered as low-quality) and remove PCR duplication reads with parameters -n 0.03 -l 20 -q 0.4 -G -d -Q 2. The coverage depth of all samples ranged from 22 to 130×.

### Long-read sequencing

For long-read DNA sequencing, 545 samples, containing 39 wild, 162 local, 117 improved, and 227 genetic stock silkworms, were selected to build Oxford Nanopore Technology (ONT) sequencing libraries. According to the standard procedure of the library construction of Oxford Nanopore Technologies Company, 20 kb *de novo* regular library of each sample was used to sequence on PromethION platform of BGI-Shenzhen. For the pooling library of two samples, reads of each sample have to be split before filter, *guppy_barcoder* v3.1.5 and *qcat* v1.0.1 were used to split reads and the intersection of two results was used as split data. The *porechop* v0.2.4 program was used to find and remove adapters. We also removed the reads with length < 5 kb and average quality less than 7. The sequencing depth ranged from 48× to 277×. The read sizes N50 ranged from 13.5 to 44.9 kb with an average of 30 kb (Supplementary Table 3).

### Transcriptome sequencing

For RNA-Seq, six tissues including cuticle, fat body, head, hemolymph, midgut and silk gland of seven samples (BomL112, BomL114, BomL122, BomL210, BomP128, BomP79, BomW44) were collected on the 3^rd^ day of the final larval stage. Total RNA was isolated using Trizol from Invitrogen Company. RNA-seq library was constructed using MGIEasy RNA Library Prep Kit and Sequenced on MGI2000 platform. All RNA sequencing was carried out at Frasergen Company (Wuhan, China). The mRNA of each tissue was collected from more than three individuals, with two biological replicates.

### SNP and Indel calling

The short reads of the 1,082 silkworm strains were mapped to the silkworm reference genome by *BWA mem* v0.7.17^36^ with default parameters using *SAMtools* v1.11^37^ and *Picard* v2.23.5 to filter the unmapped and duplicated reads. SNP and Indel were called and filtered by *GATK* v4.1.8.1^38^ (-filter “QUAL < 50.0” --filter-name LowQ - filter “DP < 200” --filter-name LowD -filter “DP > 100000” --filter-name HigD -- filter-expression “MQ < 40.0, QD < 2.0, FS > 60.0, SOR > 5.0, MQRankSum < -12.5, ReadPosRankSum < -8.0” --filter-name LowQualFilter --missing-values-evaluate-as-failing).

### Phylogenetic analysis

*VCF2Dis* v1.42 (https://github.com/BGI-shenzhen/VCF2Dis) was used to estimate *p* distances between every two samples based on VCF file containing SNPs. *PHYLIPNEW* v3.69 *fneighbor* (http://emboss.sourceforge.net/apps/cvs/embassy/phylipnew/) was used to build the Neighbor-Joining Tree. For Bootstrap Tree, SNPs were randomly selected to calculate the p distances 200 times in parallel by *VCF2Di*s (-Rand 0.25) and *PHYLIPNEW fconsense* was used to merge all the results into a consensus tree. The number of bootstrap times was changed into percentage format by *percentageboostrapTree*.*pl* in *VCF2Dis* package.

### Selective sweeps

The artificial selection regions were estimated using a sliding window approach with 5 kb window and 500 bp step size. For each window, we calculated the population divergence index (*F*_ST_), nucleotide diversity (*π*), neutrality tests (Tajima’s D), and the reduction of diversity (*ROD*) based on the prior formulas^39-41^. The top (or bottom) 1% of windows were defined as candidate selective sweep regions. The windows with selective signatures supported by at least two of the *F*_ST_, Tajima’s D, and *ROD* methods were further combined into large selection regions. In each of these regions, the genes located in or nearby (±5 kb) the extreme value window were defined as domestication or improvement associated genes. The selective sweep analysis of improvement associated regions and genes were performed in CHN-I and JPN-I, respectively.

### Genome assembly

Before assembly, *Jellyfish* v2.2.6^42^ was used to count *k*-mer frequencies (k=17). Then, the genome characters including genome size, abundance of repeat elements, and the rate of heterozygosity were predicted with *genomeScope* v1.0^43^ using NGS data.

Genome *de novo* assembly was performed with the following pipeline: (a) The ONT reads were corrected by *Canu* v1.8^44^. (b) The corrected ONT reads were assembled into contigs by *Smartdenovo* v1.0^45^ with the following parameters: wtpre – J 5000; wtzmo –k 16 –z 10 –Z 16 –U -1 –m 0.6 –A 1000; wtclp –d 3 –k 300 –m 0.6; wtlay –w 300 –s 200 –m 0.6 –r 0.95 –c 1; wtcns –m 0.6. (c) The contigs were corrected three times with ONT reads by *racon* v1.3.3^46^, and one time by *medaka* v0.7.1. In this step, *minimap2*^47^ was used to map ONT reads to the contigs, and *racon* was used to correct contigs and generated consensus sequences. The final correction was conducted using the *medaka* program. (d) The *pilon* v1.23^48^ software was used to polish the corrected contigs. In this step, NGS reads were mapped to corrected contigs using *BWA mem*, and *pilon* was used to polish with the following parameters: --fix bases --mindepth 20 --verbose --diploid. We only used the reads with mapping quality more than 20 for this step. (e) If the assembled genome exceeded the estimated size of genome survey by more than 5%, we performed de-redundancy using *Purge Haplotigs*^49^. (f) To obtain chromosome-level genomes, we used *RagTag* v1.0.0^50^ to anchor the contigs to chromosomes of the reference genome. Finally, we mapped NGS reads to the assembled genome, and assessed the genome coverage and mapping ratio. We also used Benchmarking Universal Single-copy Orthologues (*BUSCO* v 5.2.1) to estimate the integrity of the assembled genome using insecta_odb10^51^.

### Repetitive element annotation

For repetitive element annotation, simple sequence repeats were identified by *GMATA*^52^. Tandem repeats (TRs) were identified by *Tandem Repeats Finder* (TRF) v4.09^53^. Then, we used *ab initio*, structure, and homology-based methods to annotate transposable elements. Briefly, *MITE-Hunter*^54^ was used to search miniature inverted-repeat transposable elements (MITEs). *LTR_finder*^55^, *LTR_harverst*^56^, and *LTR_retriver*^57^ were used to identify long terminal repeat (LTR) retrotransposons. We performed *de novo* searching to identify repeats by *RepeatModeler* v4.1.1 (http://repeatmasker.org/), and classified the repeats into transposable elements (TEs) superfamilies through *TEclass*^58^. Finally, all TEs were merged into a repeat library which was used to annotate and mask sequence in the genome by *RepeatMasker* v4.1.1.

### Gene annotation

Three approaches including *ab initio* prediction, homology-based prediction, and reference-guided transcriptome-based assembly were used to predict genes structure in 100 genomes, including 14 wild, 41 local, 15 improved, and 30 genetic stock. For homology-based prediction, *GeMoMa*^59^ was used to align the protein sequences of the reference silkworm, *Drosophila melanogaster, Apis mellifera*, and *Danaus plexippus* to the newly assembled silkworm genomes. For the reference-guided transcriptome-based assembly, *STAR*^60^ was used to map the mRNA sequencing reads (NCBI project ID: PRJNA262539, PRJNA264587, PRJNA407019) to the newly assembled genomes, *stringtie*^61^ was used to perform RNA assembly, and *PASA*^62^ was used to predict open reading frames (ORFs). For the *ab initio* prediction, *augustus*^63^ with default parameters was used to perform *ab initio* gene prediction based on the training set that obtained by *PASA*. Finally, *EVidenceModeler* (EVM)^64^ was used to produce an integrated gene set.

Gene function information, motifs, and domains of each genome were annotated through homology searching against the public databases including Swiss-Prot, NCBI non-redundant (NR) proteins, Kyoto Encyclopedia of Genes and Genomes (KEGG), EuKaryotic Orthologous Groups (KOG), and Gene Ontology (GO). The GO items of genes were identified using the *InterProScan*^65^ program with default parameters. For the other four annotations, the proteins of each genome were used as query in *BLASTP*^66^ (e-value < 1e-5) search against Swiss-Prot, NR, KEGG, and KOG databases, respectively.

### ncRNA annotation

For non-coding RNA (ncRNA) annotation, *tRNAscan-SE*^67^ was used to identify transfer RNAs (tRNAs), *Infernal cmscan*^68^ was used to identify microRNA, rRNA and small nuclear RNA (snRNA) by searching against *Rfam* database (http://rfam.xfam.org/). Then, *RNAmmer*^69^ was used to identify rRNA and their subunits. Finally, an average of 691 (638-871) transfer RNAs (tRNAs), 126 (54-280) ribosome RNAs (rRNAs), 13 (10-18) small nuclear RNAs (snRNAs), and 6,432 (6,177-8,911) microRNAs (miRNAs) per genome were identified (Supplementary Table 4).

### Pan-gene analysis

*OrthoFinder*^70^ with default parameters was used to cluster all genes of the 100 assembled genomes into orthologous families that were classified into core, softcore, dispensable, and private gene families based on their presence frequency.

### SV identification and annotation

To identify structure variations including insertions, deletions, duplications, and inversions, the ONT reads of each sample were aligned to the reference genome using *NGMLR* v0.2.7^71^. Then, *Sniffle*s v1.0.11^71^ was used to call SVs. For filtering potentially spurious SVs, we first identified regions of the reference genome prone to producing false SV calls. To identify these regions, *SURVIVOR*^72^ was used to simulate ONT reads (∼100× the genome coverage) from the reference genome and *Sniffles* was used to call SVs. A total of 10 times simulations were performed and the 10 VCF files containing structure variations were merged by *SURVIVOR* (minimum distance = 1 kb, same SV type, and minimum SV length = 50 bp). The SVs located in these error-prone regions and their 2.5 kb flanking regions were filtered. Besides, SVs larger than 100 kb or with a “0/0” genotype were also removed. All SVs of 545 genomes were merged by *Jasmine* v1.0.1^5^ (min_support = 1, max_dist = 500, k_jaccard = 9, min_seq_id = 0.25, spec_len = 30, --run_iris).

To identify repeat elements of SV sequences, the repeat library constructed from our 100 *de novo* assembled genomes was used to annotate and mask sequences in SVs using *RepeatMasker* v4.1.1 with *RMBlast* v2.9.0-p2. The positional relationship between SV and gene was annotated using *Vcfanno*^73^.

### Estimation of pan-gene plateau

Curves describing pan-gene number were fitted with *nls* function in *R* according to previous studies^74-76^. The pan-gene number was estimated using the model *y* = *A* + *Be*^*Cx*^. To evaluate pan-gene number in groups with different number of genomes, we first selected 10 representative genomes based on phylogenetic relationships (Extended Data Fig. 1a) to carry out pan-gene number analysis. Subsequently, 10 genomes were added step by step up to 100 genomes in the following regression analyses. In total, 10 regression analyses were carried out. From the regression of 10 samples to the regression of 100 samples, the increment of gene number decreases gradually and is finally close to zero. The pan-gene increment fitted curve of each additional sample is also drawn (Extended Data Fig. 2b, c).

### Graph-based pan-genome construction

The linear reference genome and all PAVs of 545 long-read sequencing genomes were used to construct graph-based genome via *Vg toolkit*^77^. Based on this graph-based genome, we used *Vg toolkit* with the pipeline considering novel variants (used the augmented graph and gam) to call SVs in all 537 samples only sequenced through NGS technology.

### GWAS analysis based on SVs

Genome wide association studies (GWAS) were performed using *GAPIT3*^78^. Before GWAS analysis, SVs with minor allele frequency (MAF) < 0.01 were filtered. SVs phasing and imputation were performed using *SHAPEIT* v2.r837^79^. Population structure analysis was performed using *Admixture* v1.3.0^80^, the K value with the lowest cross-validation error was chosen as the best K value. The *tassel5*.*0*^81^ software was used to calculate kinship matrix. Finally, kinship matrix was used to perform GWAS analysis using multiple models containing general linear model (GLM), mixed linear model (MLM), compressed mixed linear model (CMLM), fixed and random model circulating probability unification (FarmCPU), and Bayesian-information and Linkage-disequilibrium Iteratively Nested Keyway (BLINK) implemented in *GAPIT3*.

### Identification of *Y* gene duplication

To identify the copy number of *Y* gene (carotenoid-binding protein, CBP) in each genome, the reported *CBP* transcript was used as query to perform *BLASTN*^66^ (e-value < 1E-5) against each newly assembled genome. The precise copy number and positions of *CBP* genes in each strain were obtained. Next, we performed pairwise comparison for genomic sequences of *CBP* gene in these strains with different *CBP* copy number using *BLASTN* (e-value < 1E-150) to observe their genomic structure variation. According to the results of pairwise comparison, the genome collinearity of *Y* gene region was drew using *RectChr* (https://github.com/BGI-shenzhen/RectChr/).

### Identification of large fragment variation

To identify the large fragment variations, NGS reads of selected strains were mapped to the reference genome by *BWA mem* with default parameters, or in turn, the NGS reads of reference were mapped to the assembled genome of interested strains. Then, *SAMtools* was used to calculate the coverage depth along chromosomes. *R* “*zoo*” was used to make statistics within non-overlapping 200 bp sliding windows, and the distribution of sequencing depth was drawn by *R* “*ggplot2*”. Regions without read coverage or with half (heterozygous mutant in diploid) of the average sequencing depth were regarded as PAVs and that with multiple times of the average depth were defined as duplications.

### Identification of variants underlying population traits differentiation

Considering that SVs make a greater contribution than SNPs in accounting for phenotypic variance and that high-density SNPs are more efficient compared to SVs in the identification of genomic selective signature between populations, we used a GRAF strategy that composes GWAS (genome-wide association study), ROD (reduction of diversity), AFD (allele frequencies differentiation), and *F*_ST_ (the divergence index) to identify causal variants or genes related to the interested traits.

### Identification of SV associated with strain-specific trait

We identified strain-specific SVs based on the nrSVs dataset. The specific SVs in the region of prior classical linkage analysis were defined as candidate causal SVs of the strain-specific trait. We further used *IGV*^82^ to view each bam file that generated by mapping the long reads of each strain to the reference genome and filtered the non-specific SVs.

### RNA-seq analysis and qRT-PCR assay

The RNA sequencing data of each sample was mapped to the reference genome using *bowtie2*^83^. Gene read counts were calculated using *RSEM*^84^ and gene expression was normalized using the number of Fragments Per Kilobase per Million bases (FPKM). We performed qPCR using Hieff^®^ qPCR SYBR Green Master Mix (Yeasen) reaction system on the qTOWER3G system (Analytik Jena). The primers used in qRT-PCR assay were listed in Supplementary Table 7.

### The impact of SVs on gene expression

The impact of SVs on gene expression was investigated using the previous approach^5^. Briefly, each SV-gene pair was defined based on the positional relationship between SV and its related gene. The genes that have distance less than 5 kb to the SV or contain SV were defined as SV-related genes. We filtered these SV-gene pairs to retain the pairs that had SV present in at least 2 and absent in at least 2 of the samples with RNA-seq data. For each SV-gene pair, the samples were classified into with and without SV groups. The differential expression genes between the two groups were compared using student’s *t*-test, and *p* values underwent FDR correction using the Benjamini-Hochberg procedure.

### CRISPR-Cas9 gene editing and RNA interfering

For gene knockout, the guide RNAs were designed by *CRISPRdirect*^85^, and were synthesized in BGI (Beijing, China). The Cas9 protein was purchased from Invitrogen. Then, the mixture of Cas9 protein (0.5-0.8 ng) and guide RNA (5-8 ng) was injected into newly laid eggs by microinjection.

For RNA interfering, short interfering RNAs were designed and synthesized in BGI (Beijing, China). The siRNAs (250 μM, 0.5-0.75 μl/individual) were injected into the left side of intersegment between 7^th^ and 8^th^ segment of the 3^rd^ instar larvae. Then, conductive gel was placed at the injection site (left, positive pole) and control site (right, negative pole) and 15V electric voltage was applied^86^. The sequences of sgRNA and siRNA used in this study were listed in Supplementary Table 7.

### Fineness measurement

Cocoons were reeled into single silk fibers. The silk fiber length and weight were measured. We then calculated fineness (F, dtex) of each cocoon using the formula:

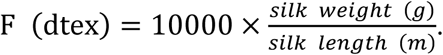

## Data and materials availability

Raw data of the long-read sequencing and short-read sequencing (including RNA-seq and whole-genome resequencing) used in this study were deposited into the CNGB Nucleotide Sequence Archive (CNSA) of China National GeneBank DataBase (CNGBdb, https://db.cngb.org), and are available through BioProject ID CNP0001815. This study also analyzes four wild silkworm genome data which available in Sequence Read Archive (SRA) database according to accession number of DRX054041, DRX054040, ERS402904, ERS402902.

## Acknowledgments

We thank Chao Su at Northwest A&F University, Bing Li at Soochow University, Keping Chen at Jiangsu University, Yuyin Chen at Zhejiang University, Weizheng Cui at Shandong Agriculture University, Chaobin Luo at Sericulture Institute of Guizhou, Yongqiang Wang at Zhejiang Academy of Agriculture Science, Wenfu Xiao, Gang Liu, and Yian Chen at Sericulture Research Institute of Sichuan, Zhanpeng Dong at Sericulture and Apicultural Research institute of Yunnan, Anjie Wang at Shandong Institute of Sericulture, Tao Fan at Sericulture Institute of Anhui, Lihui Bi at Sericulture Institute of Guangxi Zhuang Autonomous Region (GZAR), Fan Wu at Economy Crops Institute of Hubei, Yuxia Wang at Shanxi Research Institute of Sericulture, and Junwen Ai at the Sericultural Research Institute of Hunan for helping in the collection of silkworm strains. We also thank Desheng Zhang in Southwest University for ploting the figure of “Phenotypic diversity in silkworms”.

## Author contributions

F.D., W.W., X.T., and Z.X. conceived the project. F.D., Z.X., W.W., Z.T., and C.L. designed and supervised this project. F.D., H.H., X.T., S.L., Y.Y., CL.L., T.G., YH.Z., JW.L., L.Z., JH.L., W.Z., JB.S., S.H., S.W., YL.Z., Lei.Z., LL.Z., L.C., Y.T., G.C., L.Y., R.Y., H.Q., Y.L., Y.P., Y.X., T.L., and A.X. collected samples for RNA-seq and Genome sequencing, and performed phenotypic analysis. M.J.H., K.L., S.T., YC.L., S.L., JH.S., A.L., C.Z., Y.L., Z.W., W.H., J.X., T.F., Ye.Y., and J.W. performed genome assembly, SNP calling and phylogenetic analysis. M.J.H., K.L., Y.L., S.L., JH.S., A.L., and C.Z. performed genome annotation and pan-genome analysis. F.D., W.W., X.T., M.J.H., K.L., S.T., Y.L., S.L., JH.S., A.L., and C.Z. conducted the analysis of genomic variations related to complex traits. X.D., Q.G., B.Z., D.T., Y.Y., N.G., L.L., and YR.L. conducted experiments. X.T., M.J.H., K.L., S.L., JH.S., A.L., C.Z., and YC.L. interpreted data and wrote the manuscript. F.D., W.W., Z.T., E.W., and A.M. revised the manuscript.

## Extended data

**Extended data Fig. 1.**
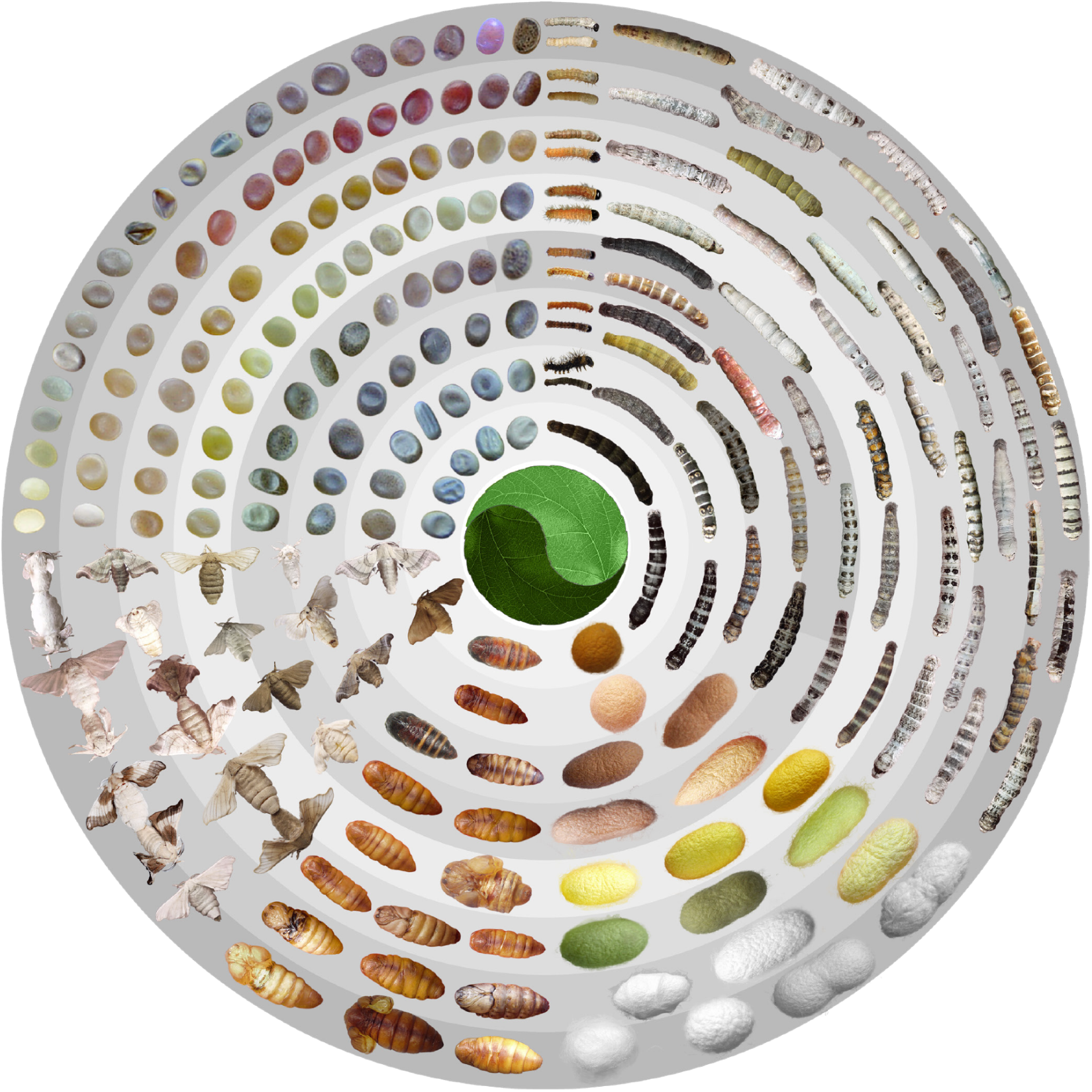
Phenotypic diversity in silkworms. Silkworms harbor enormous phenotypic diversity throughout all developmental stages from the eggs to larvae and pupae (including cocoons and pupae) and adults (clockwise from the north-west quadrant).

**Extended data Fig. 2.**
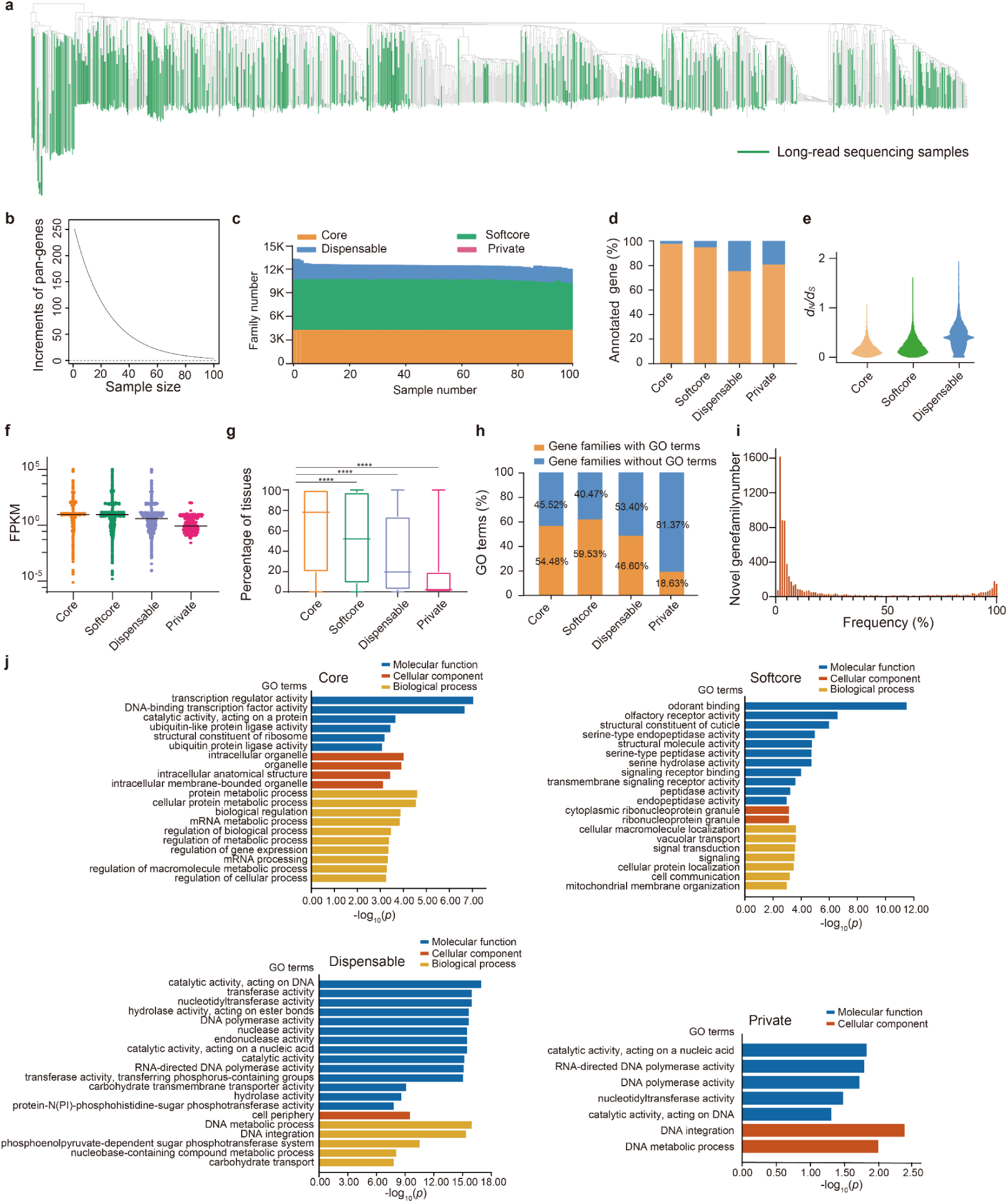
Pan-genes analysis of 100 silkworm genomes. **a**, 545 samples performed long-read sequencing are shown in green on the phylogenetic tree. **b**, Evaluation of pan-gene plateau. Derivative curve of fitted curve. When the sample size reaches 100 genomes, pan-gene increment is closed to zero. The results indicate that the pan-genes from 100 annotated genomes are representative for the species. **c**, The counts of core, softcore, dispensable, and private gene family clusters in each of the 100 genomes. The proportion of core and softcore genes (85%) is higher than that of dispensable and private genes. **d**, Proportion of genes with (orange) and without (blue) InterPro domains annotation in the four clusters. The most (98%) of core genes contain defined InterPro domains. **e**, *d*_*N*_*/d*_*S*_ values of genes in core, softcore, and dispensable groups. Core genes harbor the lowest *d*_*N*_*/d*_*S*_ values, presenting the feature of purify selection. **f** and **g**, The expression of core, softcore, dispensable, and private genes. Core genes are expressed at a higher level (f) and in more tissues (g) than dispensable and private genes. ****, *p* < 0.0001, Student’s *t*-test. **h**, The percentage of core, softcore, dispensable, and private genes with and without GO annotation. **i**, Top 20 of GO enrichment terms of core, softcore, dispensable, and private genes. Core genes are enriched in items of transcription regulator activity and DNA-binding transcription factor activity compared with the other three groups. **j**, Frequency distribution of novel genes in the100 annotated genomes.

**Extended data Fig. 3.**
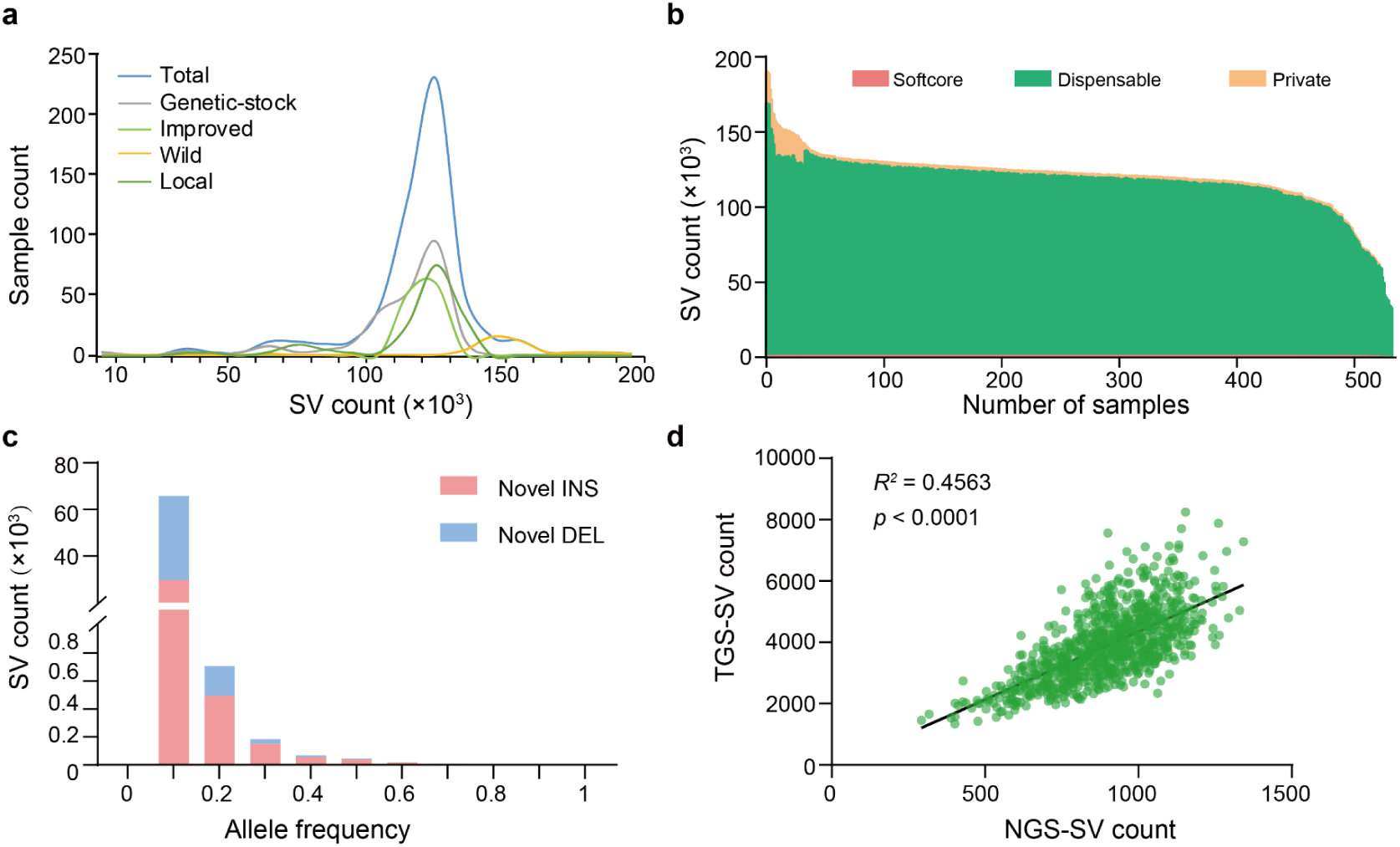
Characterization of pan-SVs in 545 silkworm genomes. **a**, Line chart shows the sample count distribution with different SV count. The SV counts are mainly distributed in 100,000-150,000 in domestic silkworms, while they are 130,000-160,000 in wild silkworms. **b**, The counts of softcore, private, and dispensable SVs for each of the 545 genomes. The dispensable SVs constitute most of all SVs in each sample. **c**, Allele frequencies of novel SVs that newly identified in 537 NGS sequenced samples by mapping short reads against the graph-based pan-genome. These novel SVs mainly appear at low frequencies among genomes. **d**, Correlations of SV count distribution along chromosomes between TGS-SV and NGS-SV. The TGS-SV was identified in the long-read sequenced genomes and the NGS-SV was identified by mapping short reads of 537 NGS sequenced samples to the graph-based pan-genome. SVs in uninterrupted 500 kb windows along chromosomes were counted.

**Extended data Fig. 4.**
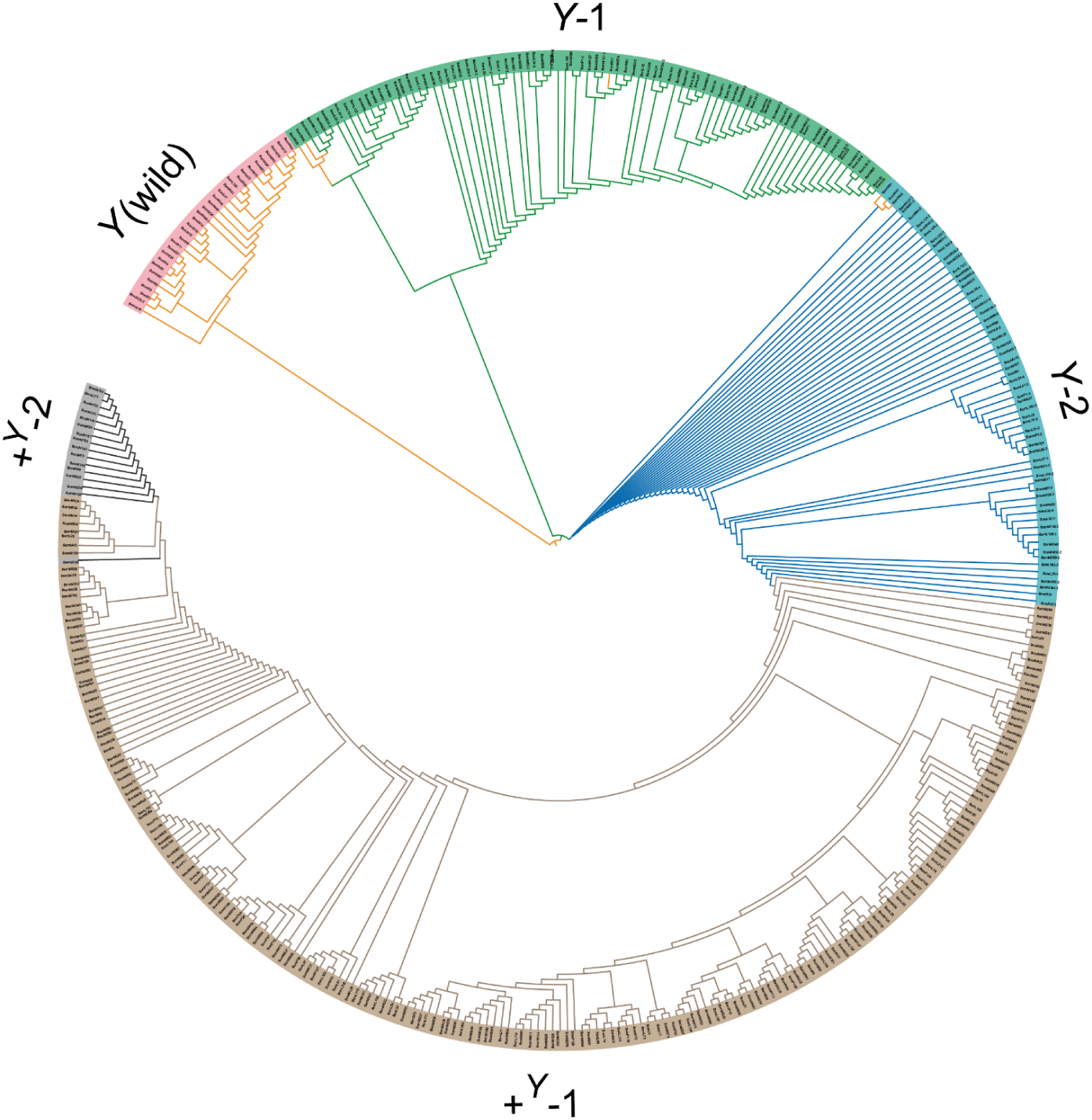
Phylogenetic tree of *CBP*. *Y*-1, *Y*-2, *+*^*Y*^-1, *+*^*Y*^-2 are the four types of *CBP* in *B. mori*.

**Extended data Fig. 5.**
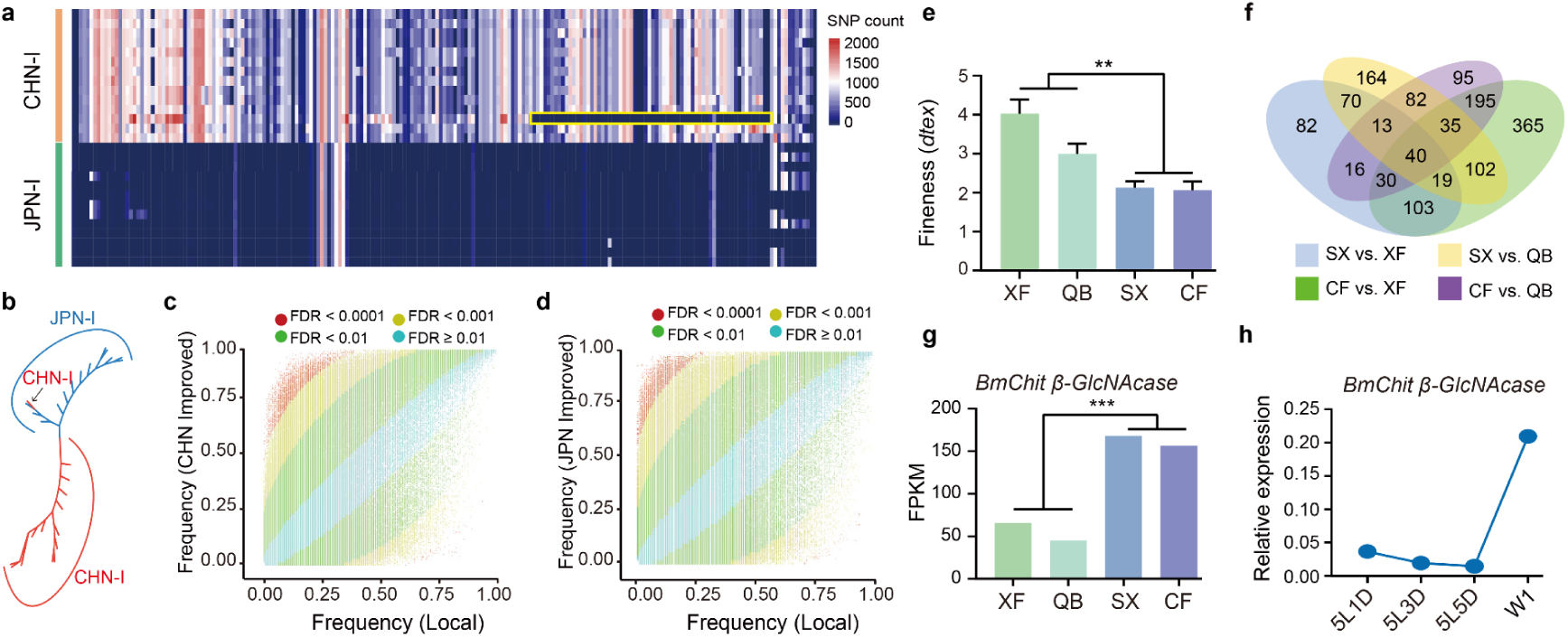
Introgression, SVs and gene related to breeding. **a**, Introgression from JPN-I (green, Chinese improved strains) to CHN-I (orange, Japanese improved strains) in chromosome 1 (yellow rectangle). Different colours on the heatmap show different SNP counts in non-overlapped 100 kb windows. **b**, The unrooted tree was constructed using SNPs in introgression region (orange rectangle) of CHN-I (red) and JPN-I (blue) and shows that one CHN-I sample (black arrow) is clustered into the JPN-I population. **c** and **d**, Frequency distribution of SVs in improvement (between local and each of CHN-I and JPN-I groups), compared with local group, 3,574 and 3,516 SVs (red dots) that present significant divergent frequencies (FDR < 0.0001, fold change > 2) were detected in CHN-I and JPN-I. **e**, The silkworm strains, CF and SX, have significantly finer silk than XF and QB strains. **, *p* < 0.01, Student’s *t*-test **f**, We performed RNA-seq of the silk press in these four strains and identified 40 differential expressed genes (DEGs). **g**, *BmChit β-GlcNAcase* was expressed significantly higher in fine silk strains (SX, CF) than in coarse silk strains (XF, QB). ***, *p* < 0.001, Student’s *t*-test **h**, *BmChit β-GlcNAcase* had an expression peak in silk press at the wandering stage (a stage at the start of spinning).

## Supplementary Table

**Supplementary Table 1**. Information and NGS sequencing summary of 1,078 samples

**Supplementary Table 2**. Nucleotide polymorphism (Pi), neutrality test (Tajima’s D), and population divergence index (*F*_ST_) in wild and domestic silkworms

**Supplementary Table 3**. Summary of ONT sequencing and structure variations of 545 genomes

**Supplementary Table 4**. Summary of annotation and pan-gene of 100 assembled genomes

**Supplementary Table 5**. Summary of annotation information of SNPs and Indels (< 50 bp)

**Supplementary Table 6**. Information of SVs associated with silkworm domestication and improvement

**Supplementary Table 7**. Nucleotide sequences of the primers, sgRNAs, and siRNAs used in experiment

